# Can exercise training improve mitochondrial thermal responses in rainbow trout cardiomyocytes?

**DOI:** 10.64898/2026.06.26.734741

**Authors:** Leteisha A. Prescott, Thuy Le, Eila Seppänen, Tiina Henttinen, Katja Anttila

## Abstract

1.

Climate-driven warming is challenging the physiological limits of aquatic ectotherms, with cardiac performance emerging as one of the key determinants of thermal tolerance. Cardiac function relies on mitochondrial ATP production, and mitochondrial dysfunction has been linked to cardiac failure at critical temperatures. However, mitochondria are plastic and may represent a target for interventions aimed at improving thermal tolerance in fish. Exercise training improves whole-animal performance in fish, including cardiac thermal performance, and improves mitochondrial function in other taxa. However, its effects on the thermal sensitivity of cardiac mitochondria remain unknown. This study investigated whether exercise-training alters cardiac mitochondrial performance at optimal and critical temperatures in rainbow trout *Oncorhynchus mykiss*. Farmed rainbow trout were subjected to a four-week exercise training regime, while control fish remained under standard rearing conditions. Cardiac mitochondrial respiration was assessed in permeabilised heart fibres at 16°C (optimal growth temperature) and 26°C (temperature associated with cardiac arrhythmia) and several biochemical and nuclear indicators were measured. No significant differences were detected between treatments for any measured variable. However, trained fish generally exhibited higher maximal respiratory capacities and respiratory control ratios, particularly at the elevated temperature, suggesting subtle improvements in mitochondrial function despite considerable inter-individual variation. Temperature influenced mitochondrial performance, increasing proton leak and reducing coupling efficiency. These findings demonstrate that cardiac mitochondrial function is thermally sensitive and represents a potential targeted for improving thermal resilience in aquaculture species.

**Summary Statement:** Exercise training did not significantly alter cardiac mitochondrial function in rainbow trout but small improvements were observed under heat stress, suggesting modest improvements in mitochondrial function despite considerable inter-individual variation.

## 3. Introduction

Anthropogenic climate change is causing significant negative impacts to aquatic environments. For aquatic ectotherms, warming waters dictate nearly all physiological processes and some species are now experiencing temperatures that surpass their thermal limits compromising health and increasing susceptibility to morbidity and mortality (Rummer et al., 2014). The teleost heart is extremely sensitive to heat stress, where changes in temperature can limit cardiac output, a key determinate of a fish’s thermal tolerance (Eliason and Anttila, 2017). When temperatures exceed an organism’s sustainable temperature zone, several molecular mechanisms are impacted, implicating cellular and organ functions, and can subsequently lead to organism death (Ekström et al., 2025; Ern et al., 2023).

Cardiac functions are highly dependent on sustained aerobic ATP production through mitochondrial oxidative phosphorylation (Brieske et al., 2025; Long et al., 2022) and therefore mitochondrial dysfunction at elevated temperatures is a concern for cardiac function. Elevated temperatures are particularly detrimental to complex I performance, causing declines in mitochondrial oxidative phosphorylation efficiency mainly because of increased proton leak (Gerber et al., 2020; Hilton et al., 2010; Iftikar and Hickey, 2013; Iftikar et al., 2015; Leo et al., 2017; Michaelsen et al., 2021; Pichaud et al., 2017). Declines in mitochondrial oxidative phosphorylation efficiency at elevated temperatures corresponds with temperatures that cause cardiac arrhythmia and is thought to be one of the key contributors to heat failure of heat-stressed fish (Hilton et al., 2010; Iftikar and Hickey, 2013; Iftikar et al., 2015).

Mitochondria are, however, a highly plastic organelle and by manipulating mitochondrial performance could provide an alternative avenue to improving organism performance. Indeed, strategies enhancing mitochondrial performance (termed mitochondrial therapies) are emerging in various industries to improve whole organism performance and health. For example, breeding, exercise training, and nutrition are all proven strategies used as mitochondrial therapies appearing in medicine (Bishop et al., 2014; Conley, 2016; Coutinho et al., 2017; Faria et al., 2023; Hsu et al., 2022; Kumar et al., 2024; Yu et al., 2023), as well as livestock (Algothmi et al., 2024; Brajkovic et al., 2023; García-Roche et al., 2022; Lapointe, 2014; Peng et al., 2021; Sanglard et al., 2023; Slaska et al., 2016; Srirattana and St John, 2017; Toyomizu et al., 2019; Wang et al., 2015) and equine (Algothmi et al., 2024; García-Roche et al., 2022; Lapointe, 2014; Peng et al., 2021) production. The role of mitochondrial therapies in aquatic production, such as finfish farming, remains elusive, despite these interventions being adaptable to commercial practices.

Exercise training could present as a promising tool for enhancing mitochondrial function in fish given moderate exercise has led to improvements in fish performance at the whole-organism level (Davison and Herbert, 2013; Espírito-Santo et al., 2026; McKenzie et al., 2020; Rodgers and Gomez Isaza, 2023a) and in other taxa, exercise has been shown to improve mitochondrial function (Bishop et al., 2014; Broome et al., 2022; Chen et al., 2021; Farhat et al., 2020; Jacobs et al., 2013; Zoladz et al., 2017). Exercise training with domesticated fish stocks enables multiple benefits and cross-tolerance abilities (Rodgers and Gomez Isaza, 2023b; Rodgers and Gomez Isaza, 2024). Exercise training enhances growth (Castro et al., 2011; Davison and Herbert, 2013; Palstra and Planas, 2011; Ytrestoyl et al., 2020), feed efficiency (Prescott et al., 2024b), athleticism (Prescott et al., 2023), disease resistance (Castro et al., 2011; Castro et al., 2013; Tierney and Farrell, 2004), immunity (Liu et al., 2018), recovery efficiency (McKenzie et al., 2012), cardiac performance (Farrell et al., 1990; Farrell et al., 1991; Nilsen et al., 2019; Zhang et al., 2016), as well as thermal (Anttila et al., 2014; Pettinau et al., 2022) and hypoxia tolerance (Fu et al., 2011). However, very few studies have attributed these exercise-enhanced phenotypes to the underlying cellular changes, such as enhanced mitochondrial function (Chen et al., 2021; Farhat et al., 2020), and as far as we know, no study has yet investigated how training could affect the thermal sensitivity of mitochondria. Exercise training could influence mitochondrial dynamics (fission-fusion), biogenesis, oxidative stress resistance, and/or ATP synthesis, all of which could enhance mitochondrial ATP production.

Equipping fish with the physiological abilities to maintain function in the face of warming waters is essential to the future and success of aquaculture and conservation restocking programs. Globally, the aquaculture industry, particularly companies farming in nearshore sea pens, shallow ponds, and/or tanks with minimal temperature control are significantly impacted by the effects of warming waters and marine heatwaves. Heat-stress mortality events are frequently reported by farmers during summer months (Broekhuizen et al., 2021; Howarth et al., 2025; IPCC, 2022), and new adaptive measures are needed to safeguard fish welfare and industry sustainability. This study evaluates exercise training as a tool to modulate mitochondrial fitness and determines if exercise-enhanced mitochondrial fitness improves mitochondrial thermal performance and thereby lends support to whole-organism thermal tolerance. By combining *in situ* functional measurements with molecular tools for abundance estimates, our study provides insight into one possible strategy to shape mitochondrial fitness in finfish.

## 4. Materials and Methods

### Ethics statement

The experiments were done according to permit administered by the National Project Authorisation Board (Permit Number ESAVI-25297-2023).

### Animal husbandry

The experimental procedures were conducted at the hatchery of the Natural Resources Institute Finland (Luke, Enonkoski, Finland) during summer 2024. Rainbow trout were initially reared in 130,000 L circular tanks and were moved to experimental tanks (1,650 L) on the 27^th^ May 2024 where they were kept under ambient temperature, oxygen, and light conditions until experiment started. Temperature and dissolved oxygen were measured daily. Water flow within the rearing tanks were maintained around 10 cm s^-1^. The fish were fed *ad libitum* with a commercial feed optimised for rainbow trout (Ova 7S, Alltech Fennoaqua, Raisio, Finland).

From the 29th – 31^st^ July, fish were randomly distributed across six experimental tanks (1,000 L); three control tanks and three exercise treatment tanks. To minimise handling stress, three fish were sacrificed to obtain an approximate fish size; fork length was 40.3 ± 0.07 cm and weight was 955 ± 1.25 g (means ± s.e.m.). The exercise treatment tanks were fitted with two pumps (Neptun NCTP-O 5000, 60W, 5000 L h^-1^, Bauhaus, Mannheim, Germany) on the inside wall next to the water inlet. The two pumps were turned on and the incoming water was fully opened for six h each weekday to generate higher flows for exercising the fish. The average flow during these periods was 40 ± 0.36 cm s^-1^ and 10 ± 0.19 cm s^-1^ for the exercise treatment and control, respectively. The exercise tanks during periods of rest exhibited flow speeds equal to the control group. Water speeds were measured twice per week for the entire four weeks of exercise training.

### Mitochondrial respiration

From the 30^th^ of August till the 8^th^ of September, mitochondrial respiration at two temperatures were measured on cardiac tissue from thirteen control and fifteen exercise trained fish. The two temperatures were 16°C and 23°C, and were chosen as they correspond to the optimal growth temperature (Jobling, 1981) for rainbow trout and temperatures that induce cardiac arrhythmia (unpublished data from this study; Pettinau et al., 2022). A control and exercise trained fish were netted out of the experimental tank and were killed with a blunt cranial blow. The fish’s fork length and weight were measured and recorded, and the heart was carefully excised, where the apex was placed in ice-cold BIOPS (2.77 mM CaK_2_EGTA, 7.23 mM K_2_EGTA, 5.77 mM Na_2_ATP, 6.56 mM MgCl_2_, 20 mM Taurine, 15 mM Na_2_phosphocreatine, 20 mM imidazole, 0.5 mM dithiothreitol, 50 mM MES, pH7.1), and two small sections were flash frozen in liquid nitrogen for later analyses.

The heart tissue submerged in BIOPS was gently teased apart using forceps to isolate fibres. Heart fibres were transferred into 2 mL ice-cold BIOPS with 20 µL of saponin on a shaker for 30 min for myocyte permeabilisation. The fibres were transferred into 2 mL of mitochondrial respiration media (MiR05; 0.5 mM EGTA, 3mM MgCl_2_, 60 mM lactobionic acid, 20 mM taurine, 10 mM KH_2_PO_4_, 20 mM HEPES, 11 mM D-Sucrose, 1g L^-1^ BSA, pH 7.1) and rinsed for 10 min. Fibres were blotted dry on a KimWipe, weighed, and ∼1 mg tissue loaded into each chamber of the O2K Oroboros oxygraphs (Innsbruck, Austria).

Two O2K Oroboros oxygraphs were used for simultaneous measurements of mitochondrial respiration at 16°C and 23°C. The oxygraph chambers were filled with 2 mL MiR05, allowed to aerate before oxygen sensors were calibrated prior to each experiment. Oxygen gas was injected into the chamber using a syringe to supersaturate the BIOPS solution (x ± S.E. = 535 ± 9 µM O_2_) and ensure oxygen did not become limited during the substrate, inhibitors, and uncouplers titration (SUIT) protocol. Respiration rates attributed to leak (non-phosphorylating) state for Complex I (Leak) were measured in the presence of 10 mM glutamate, 2 mM malate, and 15 mM pyruvate, followed by oxidative phosphorylation (CI-OXPHOS) with 12.5 mM ADP. Cytochrome C (10 µM) was added to evaluate the functional integrity of the outer mitochondrial membrane, where six fish exhibited respiration rates that increased by 15%. These fish remained in the analysis since respiration rates were only increased in one of the measurement temperatures (16°C: one control fish; 23°C: two trained and three control fish). To measure maximum phosphorylating state (CI-CII-OXPHOS) 15 mM succinate was added. Oligomycin (5 µM) was added and respiration rates were allowed to stabilise for ∼20 min, inhibiting ATP synthase (Complex V) and measuring non-phosphorylating state IV (Leak_Omy_). Carbonyl cyanide p-(trifluoro-methoxy) phenyl-hydrazone (FCCP) was titrated in steps of 2.5 µM and subsequent 0.5 µM until respiration rates plateau to uncouple respiration of Complex I and Complex II as a measure of maximum electron transport system (ETS) capacity. Rotenone (2 µM) was added to inhibit Complex I, measuring electron input through Complex II in an uncoupled state (CII). Residual oxygen consumption was measured by inhibiting Complex II with the addition of antimycin A (15 µM). Maximal capacity of complex IV was measured by raising respiration rates with ascorbate (3.2 mM) and N,N,N’,N’-tetramethyl-p-phenylenediamine dihydrochloride (TMPD; 2 mM).

Respiration rates were extracted from DatLab (DL7) as average mass-specific mitochondrial respiration rates (pmol O_2_ s^-1^ mg^-1^). Routine control ratio (RCR) was later calculated as:

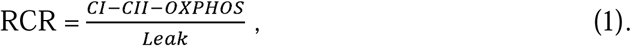

Leak control ratio (LCR) was calculated as:

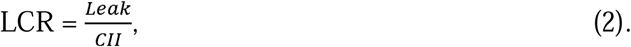

OXPHOS efficiency was calculated as:

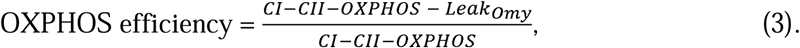

### Enzyme activity assays

Citrate synthase (CS) and lactate dehydrogenase (LDH) activities were measured from flash frozen heart samples from the same fish used to measure mitochondrial respiration. Frozen heart samples were weighed and chilled homogonisation buffer (50 mM HEPES, 1 mM EDTA, 0.1% Triton-X, pH 7.4, volume = 19 × tissue weight) were added along with two metal beads, prior to homogenising (2 min at 30 Hz) using a TissueLyser at 4°C. For CS, 15 µL of homogenate was added to 35 µL of 50 mM Tris-solution (pH 8.0) and for LDH, 7 µL of homogenate was added to 43 µL of 50 mM Tris-solution (pH 7.4). Citrate synthase activity was measured using a 384-plate containing 2.5 µL homogenate, 5 µL oxaloacetate (OA; 5mM), and 46 µL of reaction buffer (5 mM Tris, 2 mM DTNB, 12 mM Acetyl CoA, pH 8) at 412 nm and 23°C for 3 min using BioTek Synergy H1 Multimode Reader (Agilent BioTek, USA). Lactate dehydrogenase activity was measured using a 384-plate containing 2.5 µL homogenate and 24 µL NADH (0.5 mM in Tris, pH 7.4) incubated for 6 min on a plate shaker before adding 24 µL of pyruvate (50 mM in 0.5 mM of NADH) and measured at 340 nm and 23°C for 3 min using BioTek Synergy H1 Multimode Reader (Agilent BioTek, USA). CS and LDH activity were expressed as µmol enzyme mg protein^-1^ min^-1^, where protein concentration was measured from both homogenates per sample. Protein concentrations were measured spectrophotometrically with a bicinchoninic acid assay kit (ThermoFisher, Waltham, MA, USA) following manufacturers protocol.

### Mitochondrial to nuclear DNA ratio

DNA was extracted from frozen cardiac tissue samples using MACHEREY-NAGEL Genomic DNA purification kit (Thermo Scientific) following manufactures protocol. The DNA concentrations and purity were determined using a NanoDrop 200 (Thermo Scientific). For amplification of single copy nuclear gene, recombination activating 1 (Rag1), the following forward and reverse primer were used respectively: 5’-ATCCGGGTCAACACATTCCT-3’, 5’-CAATGATCCCCACATGGAG-’3. For the amplification of single copy mitochondrial gene, NADH dehydrogenase subunit 1 (mt-nd1), the following forward and reverse primer were used respectively: 5’-TAGCATACATTGTACCCGTTCTGTTAGCAG-3’, 5’-AATAGTTTTAGGCCGTCTGCGATGG-3’. The qPCR reactions were performed using 96-QuantStudioTM 12K Flex Real-Time PCR System (Thermo Scientific) with reaction volume of 20 µL with 0.5ng µL^-1^ of DNA and final primer concentrations of 0.2 µM and 10 µL of SensiFAST™ SYBR lo-ROX (Bioline). The qPCR reactions for the two genes from each sample were ran in triplicate on the same plate. DNA standards were for determination of amplification efficiency and reference sample for determining gene copy number were performed on a separate plate. The qPCR conditions were pre-amplification 95°C for 5 min, amplification 45 * (95°C for 10 s, 59°C for 10 s, 72°C for 20 s), melting curve 95°C for 5 s, 61°C for 1 min, gradual increase to 95°C for 15 s. Following methods defined in Quiros et al. (2017), mitochondrial to nuclear DNA ratio was calculated as:

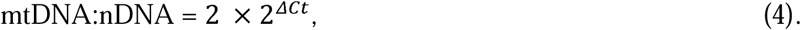

where ΔCt was calculated as:

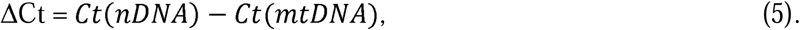

### Cardiac Energy content

Cardiac energy reserves in control and trained rainbow trout were evaluated by measuring heart glucose, glycogen, lipid, and protein concentrations.

#### Homogenisation and sample preparation

Heart tissue samples were first diluted 1:5 in 0.1 M citrate buffer (pH 5.0; citric acid monohydrate: Merck KGaA, Darmstadt, Germany; Sodium citrate dihydrate: Mallinckrodt Baker B.V., Deventer, The Netherlands) and two steel beads (2 mm diameter) were added into the sample tubes. The process was conducted on ice. The samples were then homogenised using a bullet blender (Storm24 Next Advance) at speed 8 for 4 min at 4°C. For protein and lipid assays, optimised volumes of homogenised tissue were transferred into Eppendorf tubes as follows: 5 µL for protein and 4 µl for lipid determination, and then 5 µL of citrate buffer were added to each sample. The remaining homogenate was heated at 100°C for 2 min 30s, vortexed thoroughly, and subsequently used for glucose and glycogen analyses. For each assay, 20 µL (optimised volume) of the homogenate was collected. All prepared samples were stored at -80°C for further analyses.

#### Glucose and glycogen

In the glycogen assay, 20 µL of prepared sample was incubated with 1 µL of 10 mg amyloglucosidase (A-7420, Sigma-Aldrich) in 10 mL of 0.1 M citrate buffer for 17 h at room temperature to hydrolysed glycogen into glucose. For glucose measurements, 20 µL of glucose aliquots were incubated at 4°C with the enzyme. After incubation, all samples were centrifuged at 12,000 g for 10 min at room temperature (22°C), and the supernatants were collected. For the analyses, 5 µL of supernatant and glucose standard (0, 0.35, 1, 1.5, 2, and 2.5 mg L^-1^) were mixed with 500 µL of O-toluidine reagent. The O-toluidine reagent (50 mL) was prepared by mixing 47 mL glacial acetic acid (Honeywell, Charlotte, NC, U.S.) with 3 mL O-toluidine 98% (Sigma-Aldrich, India) and 75 mg thiourea (Sigma-Aldrich, Steinheim, Germany). The reagent was stored at 4°C until use.

The samples and standard tubes were vortexed and heated at 100°C for 8 min, followed by cooling down for 4 min on ice. After vortexing, 50 µL of each sample was loaded into a 384-well plate in triplicate. Absorbance was read at 650 nm on BioTek Synergy H1 Multimode Reader (Agilent BioTek, USA). The total glycogen content was calculated by subtracting the separately measured free glucose concentration from the total glucose concentration obtained after glycogen hydrolysis and was expressed per gram of tissue.

#### Lipids and proteins

Lipids were measured by standardising against at pure olive oil solution (10mg mL^-1^; Sigma-Aldrich, Japan) dissolved in Ethanol (ETAX Aa, ethanol min^-1^. 99.5 w-%; Altia Oyj, Rajamäki, Finland). For the analysis, 4 µL of each prepared sample and 1 µL of olive oil standard (0, 0.5, 1, 2, 4, 6, 8, and 10 mg mL^-1^) were first mixed with 20 µL of H_2_SO_4_ (Merck) and heated at 100 °C for 10 min. After cooling on ice for 5 min, 1 mL of phosphovanillin reagent was added. The reagent was made by mixing 0.63 g vanillin (Sigma Chemical Company, ST. Louis, U.S.A.) with 120 mL of distilled water and 180 mL of orthophosphoric acid 13M (Thermo scientific, China). Samples and standard tubes were incubated for 15 min at 37°C, cooled down for 5 min on ice, and then 50 µL of solution were transferred to a 384-well plate in triplicate. The absorbance was read at 540 nm using BioTek Synergy H1 Multimode Reader (Agilent BioTek, USA). Lipid content was calculated based on the standard curve and expressed per gram of tissue.

Proteins were measured as previously described above using a bicinchoninic acid assay kit (ThermoFisher, Waltham, MA, USA) following manufacturers protocol.

### Statistical Analysis

All statistical analyses were performed using the R statistical language and lme4, glmmTMB, and MuMin packages. Data handling and figures were produced using tidyverse and ggplot2 packages. Model selection was assessed using Akaike’s information criterion (AIC), following selection criteria from (Richards, 2005) Model parameters (normal distribution and equal variances) were assessed using DHARMa package. Non-parametric models were used for data that failed to meet model parameters.

Normalisation of mitochondrial respiration rates was assessed in three different ways (per tissue mass, citrate synthase activity, and mtDNA:nDNA ratio), where per tissue mass was most appropriate and used for statistical analysis. For all mitochondrial respiration models, treatment and temperature were included as fixed factors and interactions between predicator and fixed variables were investigated. Repeated measures were included as a random effect nested within fish identification to account for independent responses. Fish weight, heart lipid content, and relative ventricle mass were included as co-variates and removed if not significant. Tank identification was included as a random factor, but was removed from all models during model comparisons.

## 5. Results

### Mitochondrial respiration

Mitochondrial respiration rates are presented in Fig.1, 2, and S1 and the statistical outputs are displayed in Table 1. No differences were found between treatments for any mitochondrial respiration rates. The oxygraph chamber temperature significant influenced some mitochondrial respiration states. The leak state differed significantly between temperatures with ∼4 × higher leak measured at 23°C, and similarly LCR with ∼5.4 × higher LCR at 23°C. CI-OXPHOS respiration rates differed significantly between temperatures with ∼1.3 × higher CI-OXPHOS respiration rates measured at 23°C compared to 16°C. The remaining mitochondrial respiration rates were not significantly influenced by temperature; however, leak and maximum respiration rates involving complexes I and II were close to having a significant interaction between treatment and temperature. Mitochondrial respiration rates among individuals mostly increased with elevated temperatures, although some individuals did not increase.

**Figure 1.**
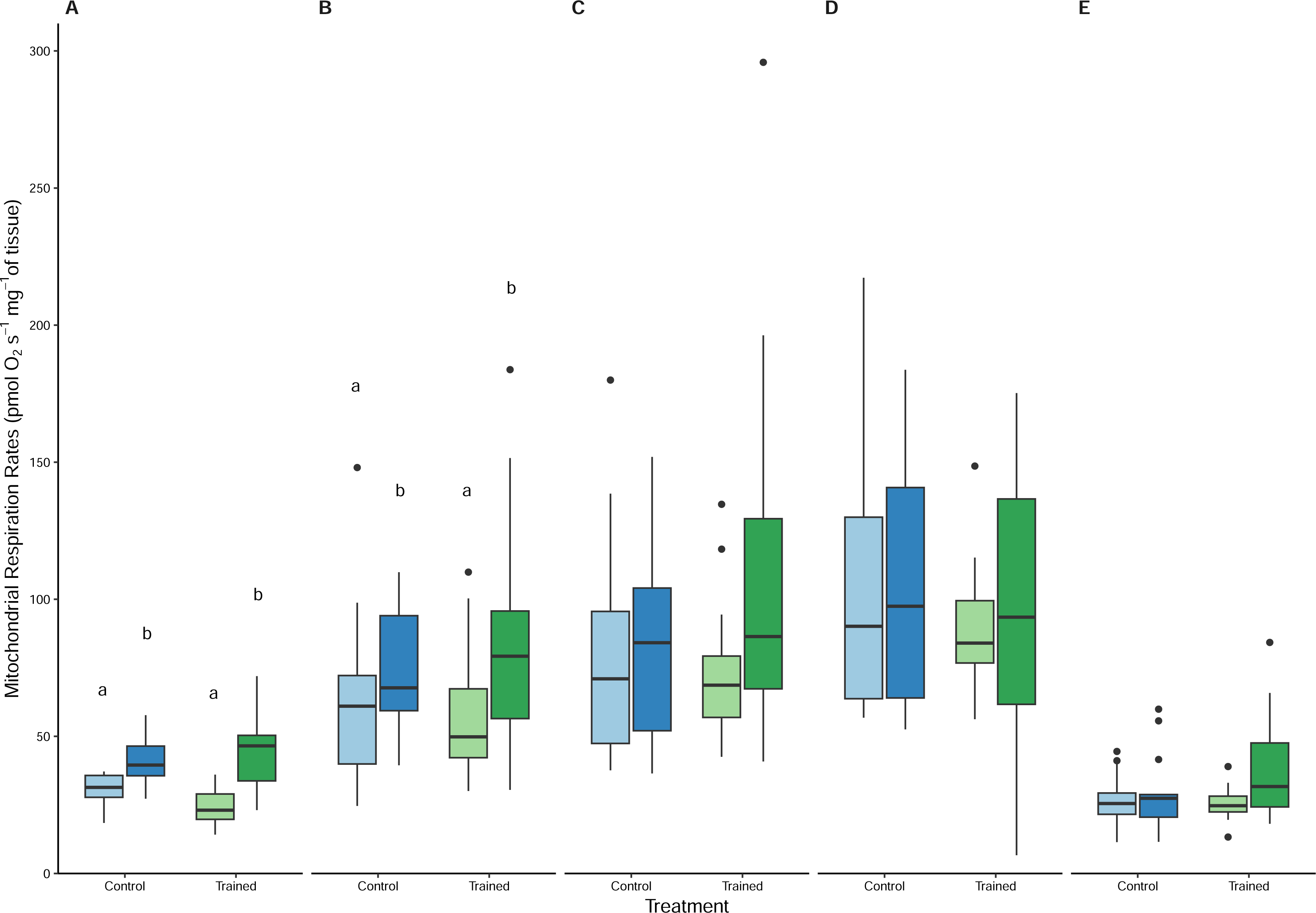
Mitochondrial respiration rates of cardiac homogenates in the presence of glutamate, pyruvate, and malate (Leak, A), ADP (CI-OXPHOS; complex I oxidative phosphorylation, B), succinate (CI-CII-OXPHOS; complex I and II oxidative phosphorylation, C), FCCP (ETS; electron transport system, D), and rotenone (CII; complex II, E) measured from control (blue; *N* = 13 individuals) and exercise trained (green; *N* = 15 individuals) rainbow trout at 16°C (lighter shade) and 23°C (darker shade). Boxplots present the median (middle bar), first and third quartiles (upper and lower bars), and the largest and smallest value within 1.5* interquartile range (IQR; vertical bars). Points represent outliers determined as values beyond the vertical bars (i.e., > third quartile +1.5*IQR, < first quartile +1.5* IQR). Letters indicate statistical differences at *P* < 0.05.

**Figure 2.**
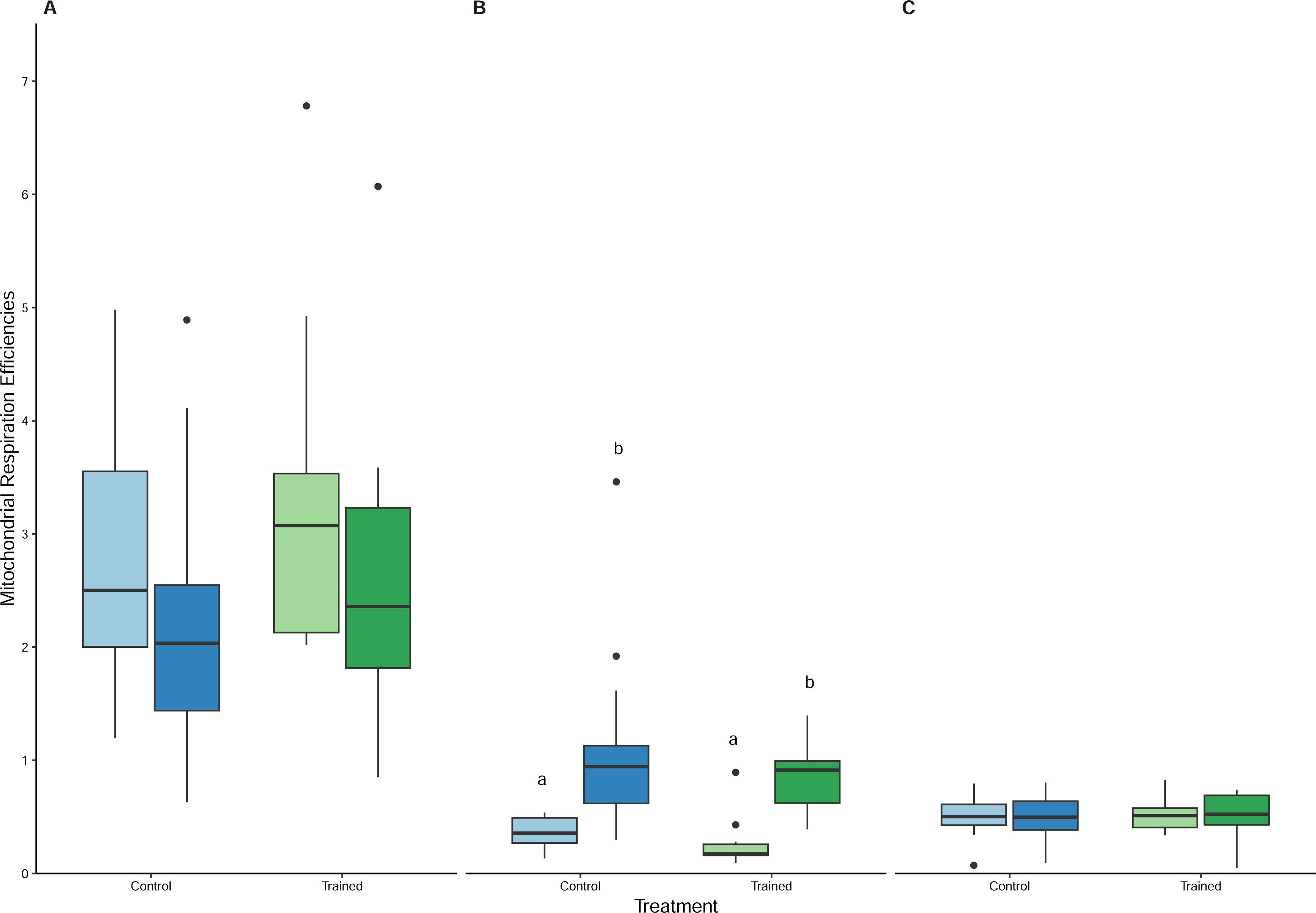
Mitochondrial respiratory control ratio (A), leak control ratio (B), and oxidative phosphorylation efficiency (C) of cardiac homogenates measured from control (blue; *N* = 13 individuals) and exercise trained (green; *N* = 15 individuals) rainbow trout at 16°C (lighter shade) and 23°C (darker shade). Boxplots present the median (middle bar), first and third quartiles (upper and lower bars), and the largest and smallest value within 1.5* interquartile range (IQR; vertical bars). Points represent outliers determined as values beyond the vertical bars (i.e., > third quartile +1.5*IQR, < first quartile +1.5* IQR). Letters indicate statistical differences at *P* < 0.05.

**Figure 3.**
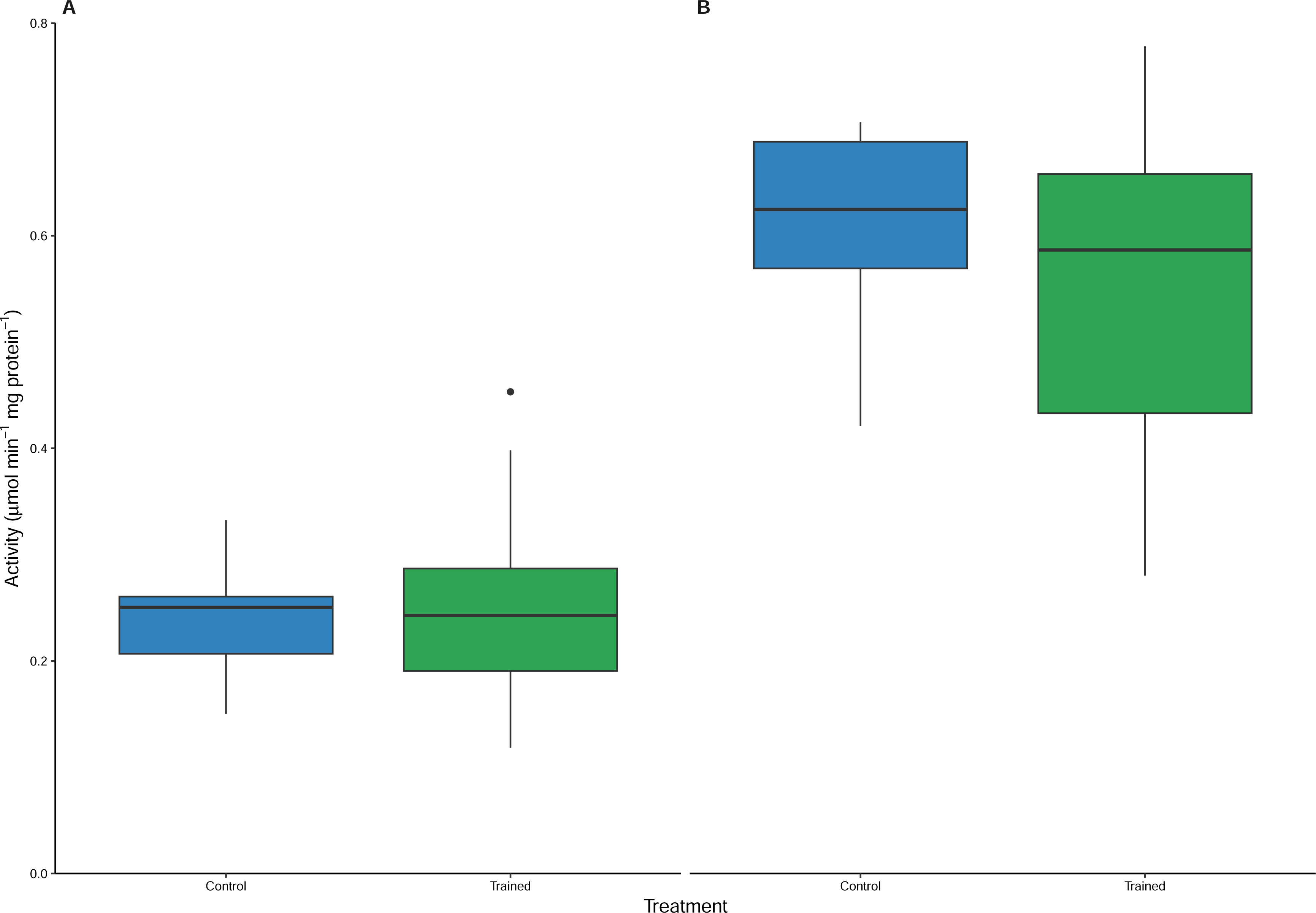
Citrate synthase (A) and lactate dehydrogenase (B) enzyme activities of cardiac tissue from control (blue; *N* = 13 individuals) and exercise trained (green; *N* = 15 individuals) rainbow trout. Boxplots present the median (middle bar), first and third quartiles (upper and lower bars), and the largest and smallest value within 1.5* interquartile range (IQR; vertical bars). Points represent outliers determined as values beyond the vertical bars (i.e., > third quartile +1.5*IQR, < first quartile +1.5* IQR).

**Figure 4.**
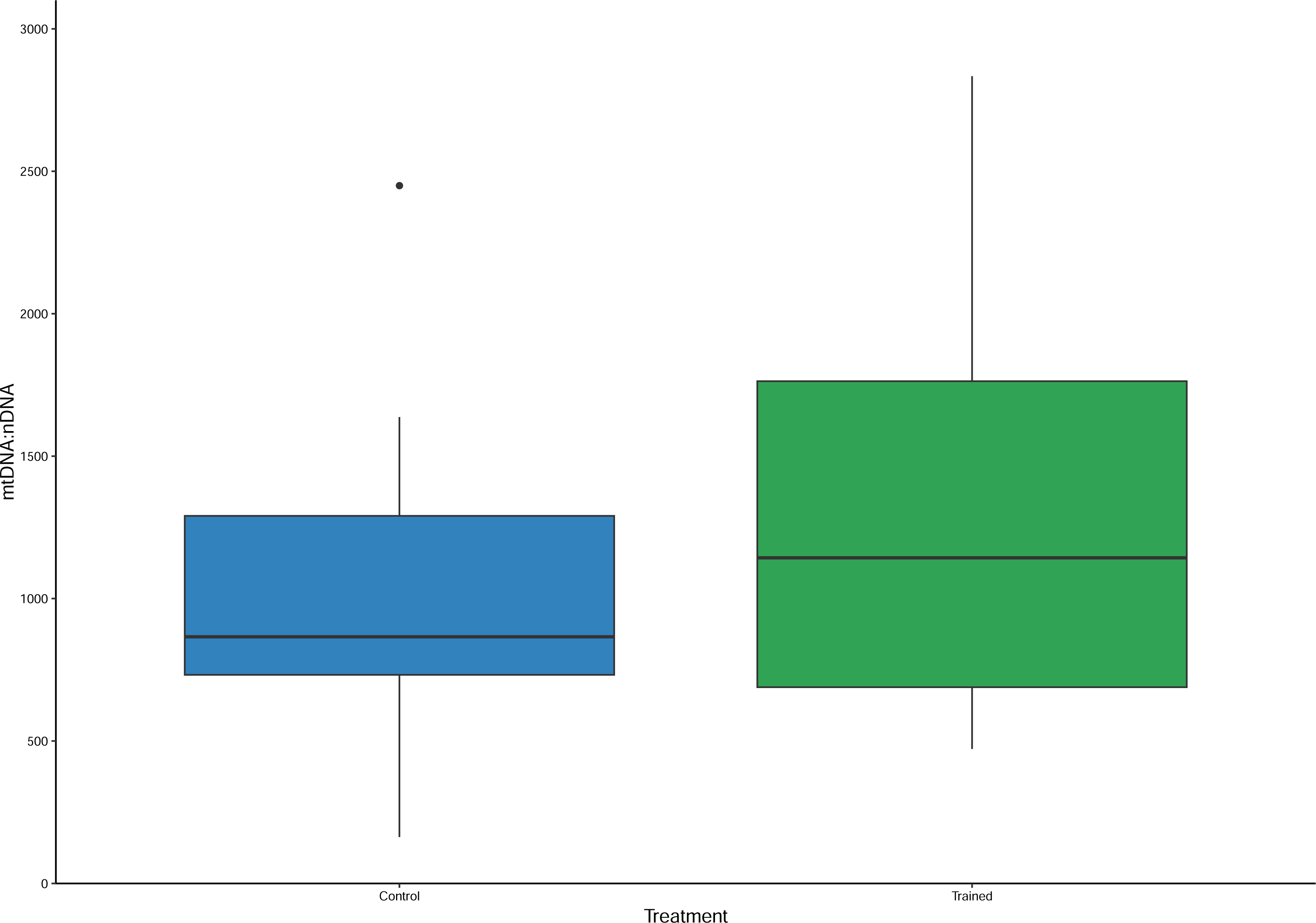
Mitochondrial to nuclear DNA ratios of cardiac tissue from control (blue; *N* = 13 individuals) and exercise trained (green; *N* = 15 individuals) rainbow trout. Boxplots present the median (middle bar), first and third quartiles (upper and lower bars), and the largest and smallest value within 1.5* interquartile range (IQR; vertical bars). Points represent outliers determined as values beyond the vertical bars (i.e., > third quartile +1.5*IQR, < first quartile +1.5* IQR).

**Figure 5.**
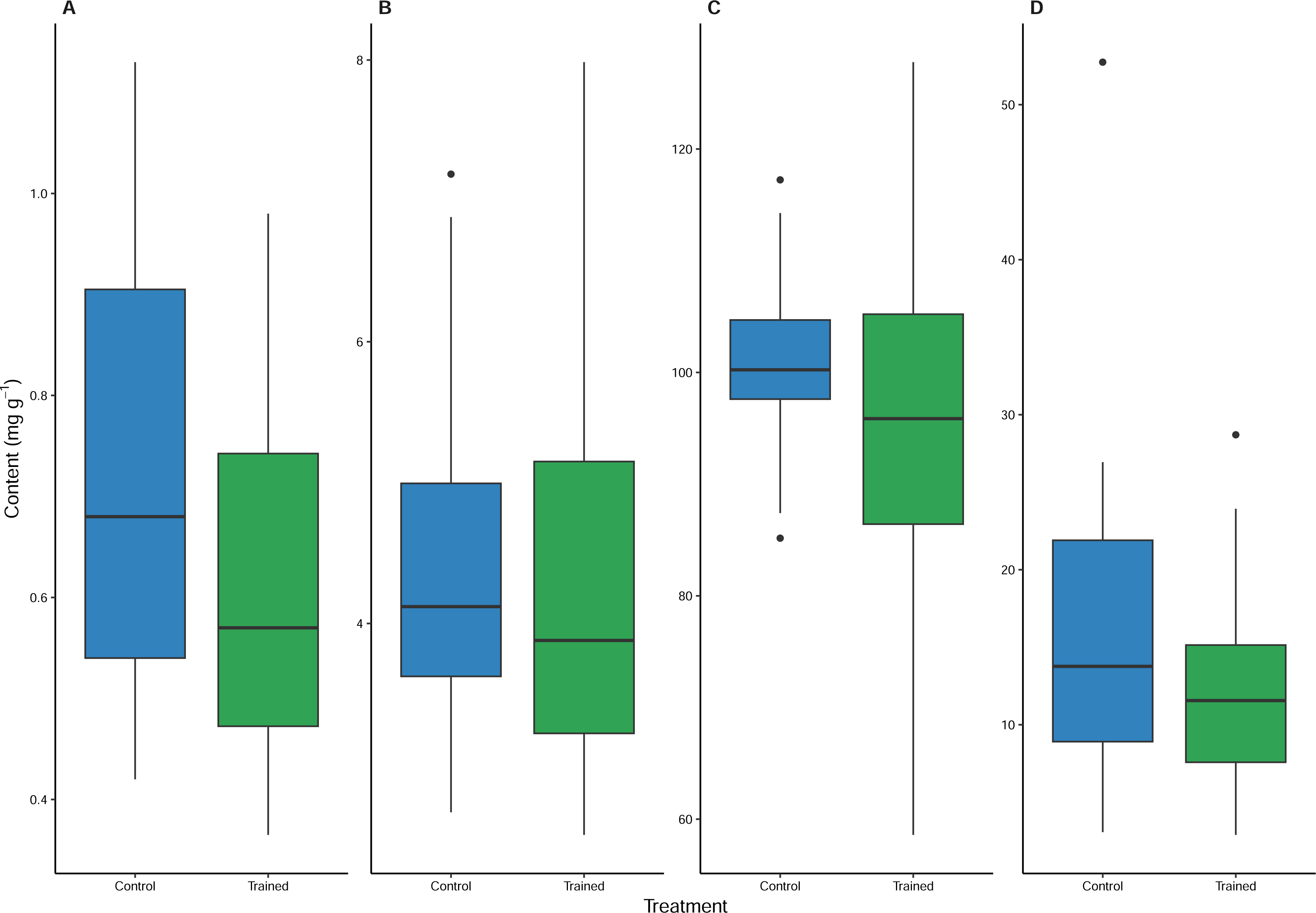
Glucose (A), glycogen (B), protein (C) and lipid (D) content of cardiac tissue from control (blue; *N* = 13 individuals) and exercise trained (green; *N* = 15 individuals) rainbow trout. Boxplots present the median (middle bar), first and third quartiles (upper and lower bars), and the largest and smallest value within 1.5* interquartile range (IQR; vertical bars). Points represent outliers determined as values beyond the vertical bars (i.e., > third quartile +1.5*IQR, < first quartile +1.5* IQR).

**Table 1.**
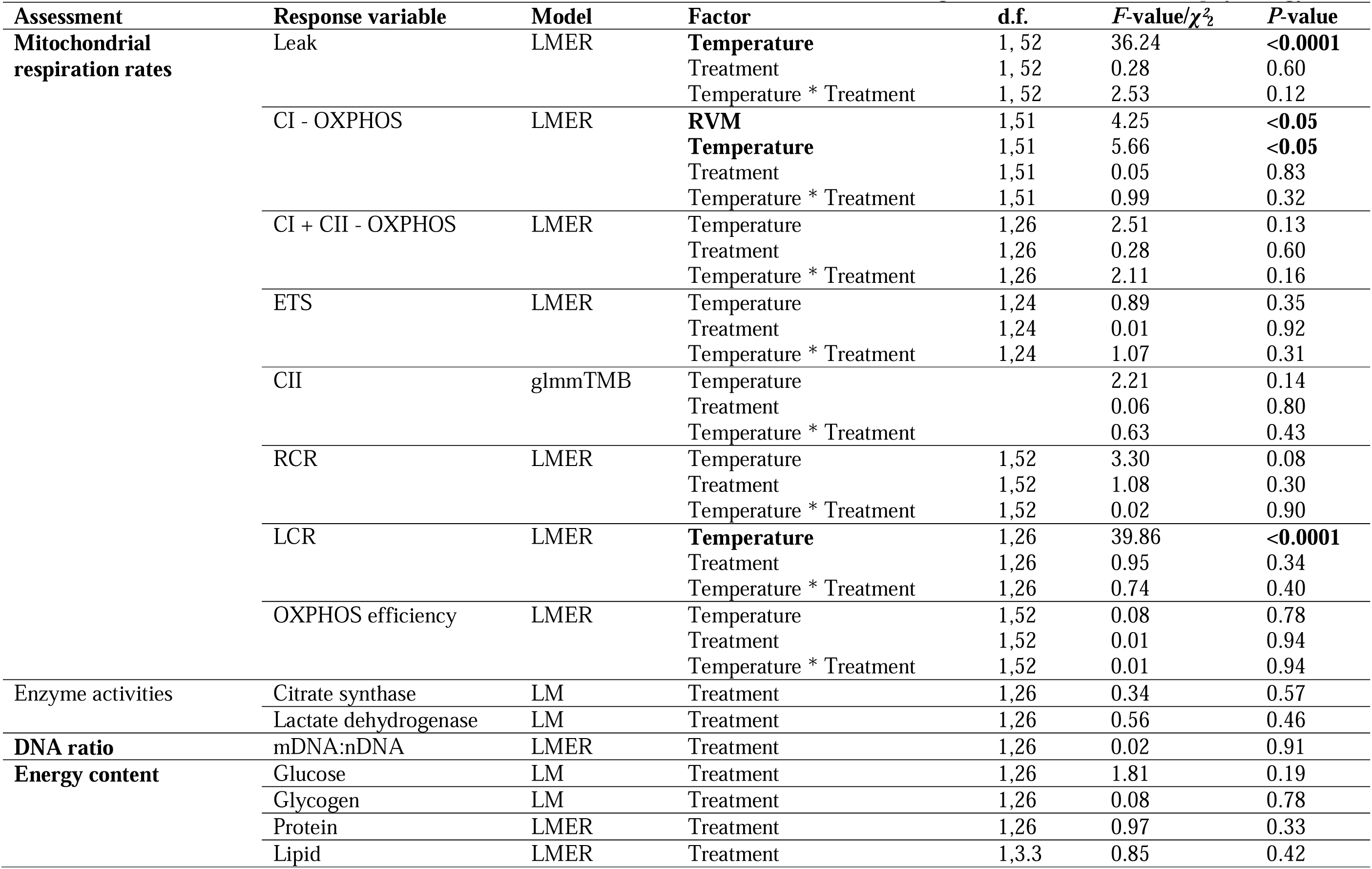
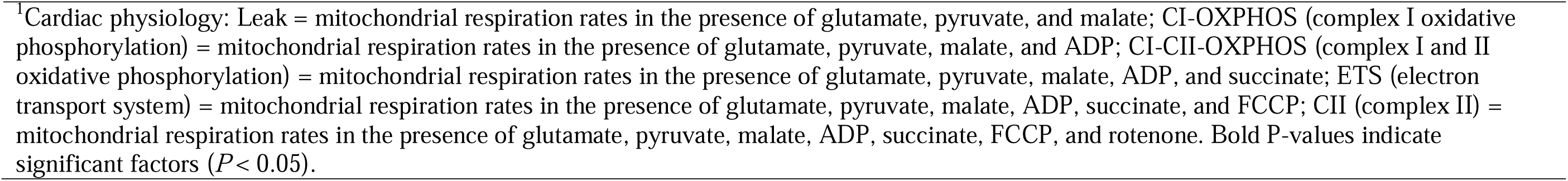
Generalised and linear mixed effects models to estimate the effects of exercise-training on rainbow trout cardiac physiology^1^.

| Assessment | Response variable | Model | Factor | d.f. | F-value/$\chi^2_2$ | P-value |
| --- | --- | --- | --- | --- | --- | --- |
| **Mitochondrial respiration rates** | Leak | LMER | **Temperature** | 1, 52 | 36.24 | **<0.0001** |
|  |  |  | Treatment | 1, 52 | 0.28 | 0.60 |
|  |  |  | Temperature * Treatment | 1, 52 | 2.53 | 0.12 |
|  | CI - OXPHOS | LMER | **RVM** | 1,51 | 4.25 | **<0.05** |
|  |  |  | **Temperature** | 1,51 | 5.66 | **<0.05** |
|  |  |  | Treatment | 1,51 | 0.05 | 0.83 |
|  |  |  | Temperature * Treatment | 1,51 | 0.99 | 0.32 |
|  | CI + CII - OXPHOS | LMER | Temperature | 1,26 | 2.51 | 0.13 |
|  |  |  | Treatment | 1,26 | 0.28 | 0.60 |
|  |  |  | Temperature * Treatment | 1,26 | 2.11 | 0.16 |
|  | ETS | LMER | Temperature | 1,24 | 0.89 | 0.35 |
|  |  |  | Treatment | 1,24 | 0.01 | 0.92 |
|  |  |  | Temperature * Treatment | 1,24 | 1.07 | 0.31 |
|  | CII | glmmTMB | Temperature |  | 2.21 | 0.14 |
|  |  |  | Treatment |  | 0.06 | 0.80 |
|  |  |  | Temperature * Treatment |  | 0.63 | 0.43 |
|  | RCR | LMER | Temperature | 1,52 | 3.30 | 0.08 |
|  |  |  | Treatment | 1,52 | 1.08 | 0.30 |
|  |  |  | Temperature * Treatment | 1,52 | 0.02 | 0.90 |
|  | LCR | LMER | **Temperature** | 1,26 | 39.86 | **<0.0001** |
|  |  |  | Treatment | 1,26 | 0.95 | 0.34 |
|  |  |  | Temperature * Treatment | 1,26 | 0.74 | 0.40 |
|  | OXPHOS efficiency | LMER | Temperature | 1,52 | 0.08 | 0.78 |
|  |  |  | Treatment | 1,52 | 0.01 | 0.94 |
|  |  |  | Temperature * Treatment | 1,52 | 0.01 | 0.94 |
| Enzyme activities | Citrate synthase | LM | Treatment | 1,26 | 0.34 | 0.57 |
|  | Lactate dehydrogenase | LM | Treatment | 1,26 | 0.56 | 0.46 |
| **DNA ratio** | mDNA:nDNA | LMER | Treatment | 1,26 | 0.02 | 0.91 |
| **Energy content** | Glucose | LM | Treatment | 1,26 | 1.81 | 0.19 |
|  | Glycogen | LM | Treatment | 1,26 | 0.08 | 0.78 |
|  | Protein | LMER | Treatment | 1,26 | 0.97 | 0.33 |
|  | Lipid | LMER | Treatment | 1,3.3 | 0.85 | 0.42 |

<sup>1</sup>Cardiac physiology: Leak = mitochondrial respiration rates in the presence of glutamate, pyruvate, and malate; CI-OXPHOS (complex I oxidative phosphorylation) = mitochondrial respiration rates in the presence of glutamate, pyruvate, malate, and ADP; CI-CII-OXPHOS (complex I and II oxidative phosphorylation) = mitochondrial respiration rates in the presence of glutamate, pyruvate, malate, ADP, and succinate; ETS (electron transport system) = mitochondrial respiration rates in the presence of glutamate, pyruvate, malate, ADP, succinate, and FCCP; CII (complex II) = mitochondrial respiration rates in the presence of glutamate, pyruvate, malate, ADP, succinate, FCCP, and rotenone. Bold P-values indicate significant factors ( $P < 0.05$ ).

### Enzyme activity, mitochondrial to nuclear DNA ratio, and energy content

Exercise training did not influence cardiac enzyme activities, as citrate synthase and lactate dehydrogenase did not differ between treatments. The mitochondrial to nuclear DNA ratio did not differ between treatments either. Cardiac energy content was also not influence by exercise training with glucose, glycogen, protein, and lipid contents not differing between treatments.

## 6. Discussion

Cardiac mitochondrial oxidative phosphorylation is a key contributor to cardiac dysfunction during thermal stress and therefore presents a focal point for better understanding, predicting, and enhancing organism thermal tolerance. For finfish farmers, applying this understanding of how mitochondrial function underpins organism performance and stressor tolerance could enable identification of new strategies to improve mitochondrial function and stock performance. We examined how exercise training alters cardiac mitochondrial function in rainbow trout at temperatures associated with optimal growth and cardiac arrhythmia.

Exercise training resulted in modest changes in mitochondrial function, with some effects like leak, CI-CII-OXPHOS, and maximum respiration rates, becoming more apparent at critical temperatures. Furthermore, trained fish appeared to be more responsive to temperature at the higher temperature, although considerable inter-individual variation was observed. Temperature strongly influenced mitochondrial performance, with complexes I and II appearing particularly sensitive, suggesting a limitation for ATP production under thermal stress.

### Exercise-enhanced mitochondrial function

Exercise training in mammalian species improves mitochondrial ATP production through modulating fusion and fission processes, such as changes in mitochondrial abundance, morphology, and efficiency (Djalalvandi and Scorrano, 2022; Li et al., 2025). In the present study, exercise training induced modest, non-significant changes to mitochondrial dynamics, where small increases in mtDNA:nDNA and respiratory efficiencies were found and little change in citrate synthase activity, suggesting limited mitochondrial remodelling under these conditions. However, exercise-trained fish in general exhibit higher maximal respiratory capacity, involving complex I and II, with differences most evident at critical temperatures. This pattern, alongside elevated respiratory and leak control ratios at both temperatures indicates that exercise may enhance mitochondrial efficiency. However, absolute leak was only lower than control fish at optimal temperatures and were slightly higher under challenging temperatures. These mitochondrial responses are somewhat consistent with the small, non-significant increases in critical thermal maximum (CT_max_) measured from a subsample of rainbow trout held under the same conditions in an analogous study and improvements in whole animal cardiac thermal tolerance observed in a previous study (Pettinau, 2023; Pettinau et al., 2022). Together, these small modifications suggest exercise training can make modest contributions to mitochondrial function to improve thermal tolerance, but further exploration is needed.

Improved mitochondrial thermal responses driven by exercise training in rainbow trout may have been masked by substantial inter-individual variability within treatments. High variability was observed across mitochondrial respiration rates in both treatments, but particularly among trained fish when measured under elevated temperatures. This suggests that some individuals may respond more favourably to exercise-training than others, while for some fish the training regime itself may represent an additional physiological stressor or oppositely, not energy demanding enough. Contrasting responses to exercise-training may also reflect domestication pressures and genotype-by-environmental interactions (Prescott et al., 2024a). These individual responses are illustrated in Fig. S1, where some individuals (e.g., 8/13 control and 9/15 exercise trained individuals increased complexes I and II maximum mitochondrial respiration rates with elevated temperatures) exhibited increased respiration rates with elevated temperatures, whereas others showed marked declines. This pattern suggests that certain individuals were able to maintain mitochondrial function under challenging temperatures, while others exhibited signs of mitochondrial dysfunction. Future research should investigate whether individuals with divergent mitochondrial thermal responses differ in thermal tolerance and whether these traits are associated with responsiveness to exercise training. This would enable better understanding of the drivers of individual physiological variation and define reliable strategies for improving production stock’s thermal performance.

Although there were only some links evident between exercise training influencing mitochondrial ATP production, exercise training remains a promising strategy to enhance cardiac performance and thermal resilience in vulnerable stocks. Previous studies using similar training regimes have reported improvements in cardiovascular and thermal performance in rainbow trout, although these were conducted in smaller individuals (Papadopoulou et al., 2022; Pettinau et al., 2022). In the present study, cardiac energy content did not differ significantly between groups; although reductions in glucose and lipid levels in the exercise-trained fish suggest that the exercise-training regime imposed a greater energetic demand. This indicates that the training stimulus may have been sufficient to alter substrate utilisation, but insufficient to drive substantial mitochondrial remodelling.

In other species, exercise training lowers the resting heart rate and overall energy demand during exercise (Hellsten and Nyberg, 2015). If fish also exhibit these cardiovascular changes with exercise training, it could also provide explanation why significant modulations in mitochondrial respiration rates weren’t observed and that slight changes in mitochondrial content was sufficient to meet the energy demands of increased energy expenditure. As such, more intensive exercise regimes requiring larger energy demands may have elicited greater changes in mitochondrial dynamics. Further optimisation in training intensity and duration may therefore be necessary to elicit more pronounced physiological and performance benefits, and the use of a varying flow regime could accommodate for individual variability.

Nonetheless, the modest changes in mitochondrial function observed provide some support for the altered performance observed in previous studies in exercise-trained fish (Anttila et al., 2014; Farrell et al., 1990; Palstra et al., 2015; Pettinau et al., 2022; Timmerhaus et al., 2021), suggesting that even small shifts in mitochondrial performance may influence cardiac function under thermal stress (Pettinau et al., 2022). Beyond mitochondrial efficiency, additional cellular mechanisms may contribute to exercise-induced changes in thermal performance. For example, exercise training has been shown to modify the electrical excitability of cardiomyocytes in rainbow trout, including prolongation of the action potential (Le et al., 2026), which could influence cardiac stability at higher temperatures. Structural adaptations may also play a role; in mammals, exercise increases mitochondrial cristae density, enhancing oxidative phosphorylation capacity without requiring large increases in mitochondrial volume (Djalalvandi and Scorrano, 2022; Li et al., 2025; Nielsen et al., 2017). Furthermore, improved myocardial perfusion through increased capillary density and cardiac remodelling has been document in humans (Pinckard et al., 2019), potentially supporting oxygen delivery and metabolic demand during stress. Together, these mechanisms suggest that exercise-induced improvements in thermal performance may arise from integrated changes across multiple levels of cardiac organisation, rather than mitochondrial function alone.

### Mitochondrial function thermal sensitivity

Temperature had a strong influence on mitochondrial respiratory performance. Elevated temperatures increased leak respiration and the leak control ratio, indicating reduced mitochondrial membrane integrity and increased proton permeability, which together lowers OXPHOS efficiency. Although CI-linked respiration rates were higher at elevated temperatures, this likely reflects accelerated metabolic flux rather than improved efficiency, as increased oxygen consumption was accompanied by greater uncoupling. A similar, albeit not significant, trend was observed for CII respiration. These responses are consistent with previous studies showing that elevated temperatures disproportionately impair complex I function and increase proton leak, ultimately reducing mitochondrial efficiency. This has been demonstrated in triplefin species (Family Tripterygiidae; Hilton et al., 2010), spotty wrasse *Notolabrus celidotus* (Iftikar and Hickey, 2013; Iftikar et al., 2015), rainbow trout *Oncorhynchus mykiss* (Michaelsen et al., 2021; Pichaud et al., 2017), Atlantic salmon *Salmo salar* (Gerber et al., 2020), Polar cod *Boreogadus saida* (Leo et al., 2017), and Atlantic cod *Gadus morhua* (Leo et al., 2017). The reduced mitochondrial ATP production efficiency observed here occurred at temperatures associated with cardiac arrhythmia in rainbow trout, supporting the idea that mitochondrial dysfunction contributes to cardiac failure under heat stress. Together these findings highlight a key trade-off at elevated temperatures, where increased respiratory rates are offset by reduced coupling efficiency, ultimately limiting ATP supply to the heart.

### Conclusions and implications for industry

Overall, our findings demonstrate that mitochondrial function plays a central role in shaping cardiac performance under thermal stress, with elevated temperatures driving increased respiratory demand but reduced coupling efficiency, ultimately limiting ATP supply. While exercise training elicited only modest improvements in mitochondrial performance, the enhanced responses observed under critical conditions suggest that mitochondrial function retains some plasticity and may be a viable target for intervention. In particular, the sensitivity of complex I-linked respiration to temperature highlights a potential bottleneck in cardiac energy metabolism during heat stress. For aquaculture, these findings suggest that strategies aimed at improving mitochondrial efficiency may enhance thermal resilience in farmed fish. However, the limited magnitude of responses observed here indicates that optimisation of such approaches will be necessary to achieve commercially relevant gains in performance. In particular, how large individual variance can drive population outcomes where some fish respond well to the training regime used and others did not. Perhaps, a varying flow regime that allows fish to select their swimming speed and thus effort could be a viable approach for industry. Future work should therefore focus on identifying interventions that more effectively target mitochondrial function and its integration with whole-organism physiology under warming conditions.

## Acknowledgements

The authors thank the staff of LUKE aquaculture facilities in Enonkoski, Finland for efforts towards fish husbandry and assistance throughout the trial, Tytti Uurasmaa and staff of the Department of Biology, University of Turku for technical assistance.

## Competing Interests

The authors declare no competing or financial interest.

## Funding

The authors acknowledge the financial support of the Blue Economy Cooperative Research Centre (CRC), established and supported under the Australian Government’s CRC Program, grant number CRCXX000001 (previously 20180101). The CRC Program supports industry-led collaborations between industry, researchers, and the community. The authors acknowledge financial support from the Finnish National Agency for Education, The Company of Biologists, and the Australian New Zealand Marine Biotechnology Society. The authors acknowledge the financial support from the Academy of Finland (Project 350315).

## Data availability

Data will become available upon request.

## Authors’ contributions

Conceptualisation: all authors; Methodology: all authors; Formal analysis: L. A. Prescott; Writing: L. A. Prescott wrote original draft, and all authors were involved in reviewing and editing the manuscript; Funding acquisition: L. A. Prescott and K. Anttila.

## Notes

### Competing Interest Statement

The authors have declared no competing interest.

