## Supplementary Material for "Can exercise training improve mitochondrial thermal responses in rainbow trout cardiomyocytes?"

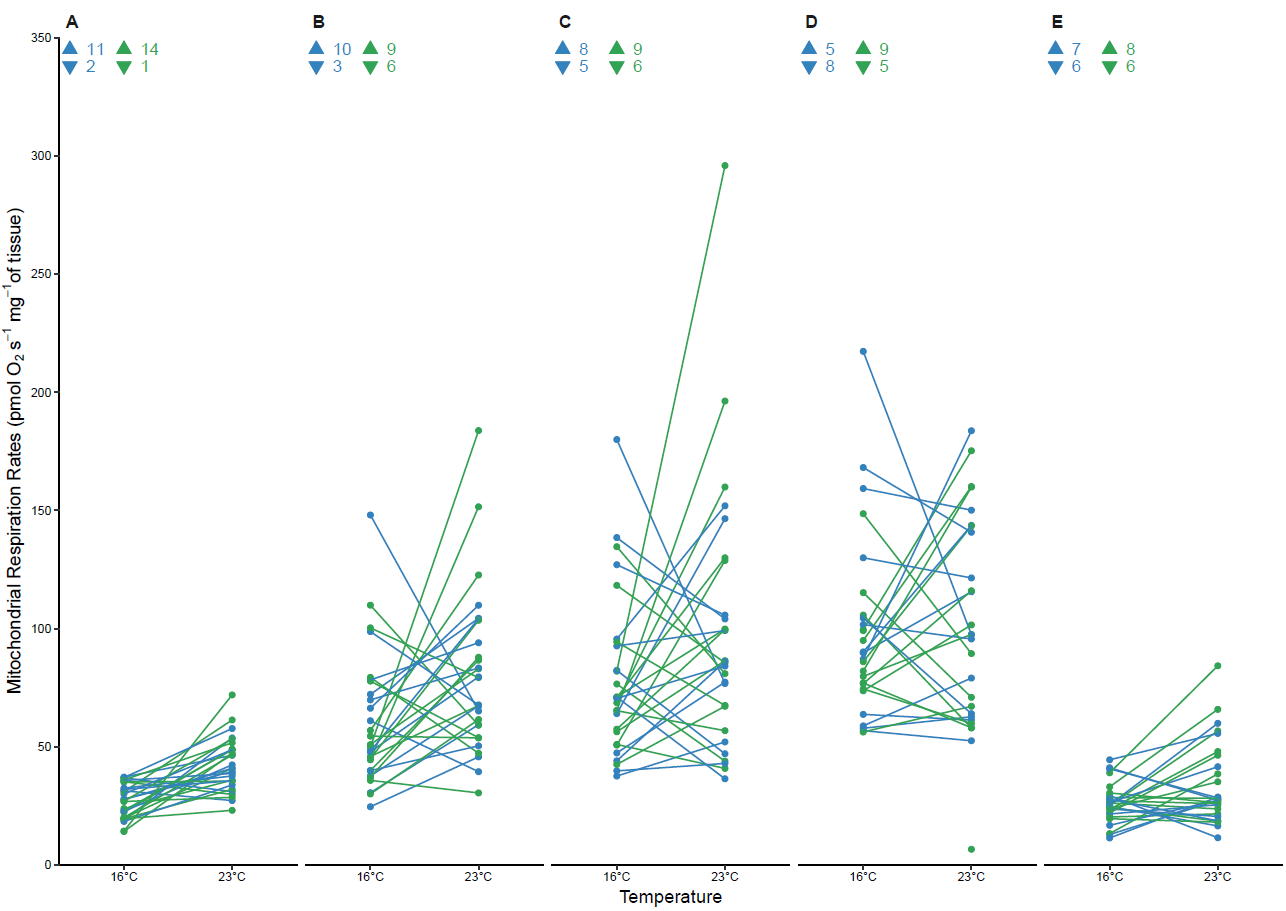

Fig S1. **Individual mitochondrial respiration rates of cardiac homogenates in the presence of glutamate, pyruvate, and malate (Leak, A), ADP (CI-OXPHOS; complex I oxidative phosphorylation, B), succinate (CI-CII-OXPHOS; complex I and II oxidative phosphorylation, C), FCCP (ETS; electron transport system, D), and rotenone (CII; complex II, E) measured from control (blue; *N* = 13 individuals) and exercise trained (green; *N* = 15 individuals) rainbow trout at 16°C (lighter shade) and 23°C (darker shade).** Points and lines represent individual fish. Blue (control) and green (exercise trained) upward and downward arrows indicate the number of individuals that increased or decreased mitochondrial respiration rates with an increase in temperature.
